# Evidence of selection in sympatry between two closely related species of rockfish

**DOI:** 10.64898/2026.09.15.751943

**Authors:** SP Harned, BD Scholten, MO Burford Reiskind

## Abstract

Ecological or reproductive barriers can maintain species boundaries by preventing introgression in closely related taxa when their distributions overlap (sympatry). In sympatry, sister-taxa may have greater genetic divergence compared to allopatric parts of their range. When analyzing populations of one of the sister-taxa, this divergence may translate to greater genetic differentiation between sympatric and allopatric populations of this species’ range. This genetic differentiation can be caused by either genetic drift or natural selection, depending on the evolutionary history of secondary contact. To identify a selective process, it is critical to find genes responsible for maintaining species barriers in sympatry. Here, the role of natural selection in genetic differentiation was examined within two recently diverged rockfish species, *Sebastes diaconus* and *Sebastes mystinus*. These species overlap along over 400 km of coastline in the eastern Pacific, with no evidence of hybridization in this sympatric region. This study found evidence of geographic genetic differentiation across a large span of the *S. diaconus* range, but not within the *S. mystinus* range. For both species, outlier loci were identified that were associated with regions of the genome under directional selection in allopatric versus sympatric populations. This study also found signals of directional selection in shared genomic regions of both species, including in genes involved in neurological function, photoreception, and gene regulation. This suggests that the evolutionary process of reinforcement maintains species boundaries when the two species come into secondary contact.

## Introduction

To understand the process of speciation requires identifying genes that maintain species boundaries in recently diverged species in secondary contact, the process by which two previously allopatric species overlap again. Species boundaries between species in secondary contact may result from genetic drift or natural selection, depending on the species’ evolutionary history and timing of secondary contact between the sister-taxa. If the speciation process was not complete before secondary contact, genetic analysis of one of the two species throughout its range may reveal different degrees of genetic differentiation when comparing sympatric populations to allopatric populations (where this species do and do not co-occur with their sister-taxa, respectively). Previous studies show that within overlapping geographic distributions of sister-taxa, there is often reproductive isolation caused by physical or phenotypic barriers that maintain genetic separation between species and prevent widespread introgression, termed evolutionary reinforcement. Yet these same barriers may not exist in allopatric populations (e.g. ^1–4^). Ecological segregation, such as habitat specialization or depth delineation, creates a physical barrier that reinforces species barriers, and this niche partitioning is broadly referred to as character displacement.^5,6^ Previous studies have found character displacement in many species, including stickleback fishes, spadefoot toads, and anole lizards.^6–8^ One mechanism for character displacement is directional selection of genes responsible for mating behavior differences, referred to as prezygotic reproductive barriers.^9,7,8^ In other cases, reproductive character displacement may be driven by lower fitness of the hybrids causing post-zygotic reproductive barriers.^10–14^ Distinguishing among different scenarios of reproductive reinforcement during secondary contact between two recently diverged species presents a challenge in the study of the speciation process. Determining which evolutionary force (genetic drift or natural selection) or isolating mechanism is the driving force for the two sister-taxa will reveal more about the process of speciation.

There are many factors that contribute to evolutionary reinforcement, including ecological segregation or niche partitioning, random changes in isolating barriers due to genetic drift, selection on genes associated with prezygotic or postzygotic reproductive barriers, postzygotic reproductive barriers, or pleiotropic effects on mating fitness. First, it is critical to measure the degree of genetic differentiation between the two sister-taxa throughout their geographic range in both allopatric and sympatric zones. For example, if speciation was complete prior to secondary contact, we would predict there would be the same degree of genetic divergence between the species when we compared allopatric or sympatric populations and there is no evidence of introgression. Alternatively, if speciation was not completed prior to secondary contact, we predict we would find evidence of selection in sympatric compared to allopatric populations within a species range and/or evidence of introgression between the two species in sympatry. Depending on the degree of genetic divergence in sympatry and/or introgression we can determine whether we are early in the evolutionary reinforcement process or they are not distinct species. If we find the former without the latter, meaning they do not introgress in sympatry, an established reinforcing mechanism is likely in place and likely maintaining species boundaries. This means we are closer to the completion of the speciation process via evolutionary reinforcement (Fig. 1). To confirm evolutionary reinforcement would require a thorough understanding of the evolutionary history of both species, elevated divergence in the sympatric zone, low to no introgression in sympatry, and signatures of selection within the genome in the sympatric zone.

**Fig 1.**
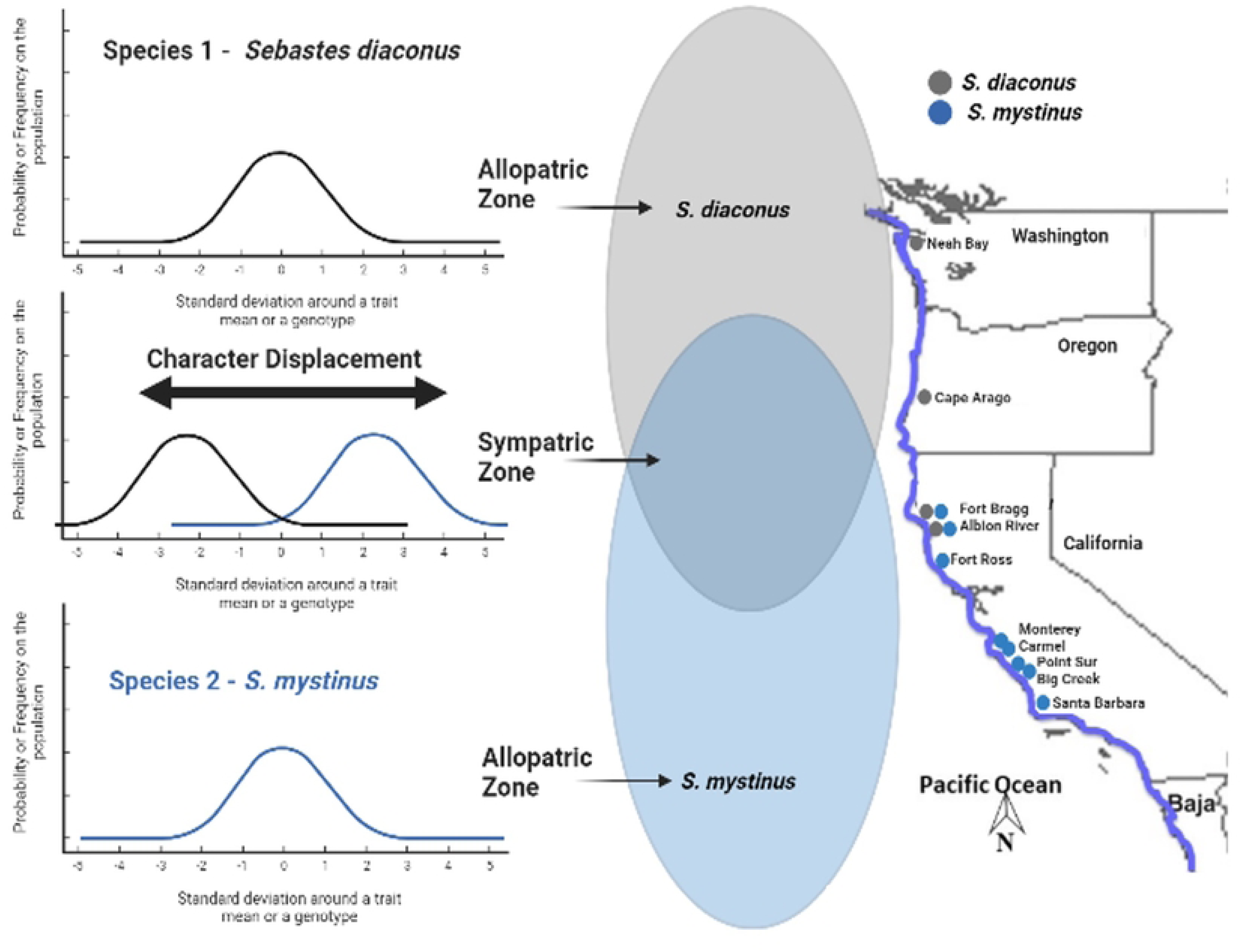
Character displacement as a mechanism maintaining species boundaries between *Sebastes diaconus* and *S. mystinus* in sympatry. Map to the right indicates the sampling sites. The nine individuals (Table S1) found in the southern group of *S. diaconus* were sampled from at or near Fort Ross, Monterey, Point Sur & Big Creek. Image was created in biorender.com.

The *Sebastes* genus underwent a historic evolutionary radiation event approximately 5 million years ago, giving rise to over 60 species and subspecies.^15,16^ Two previous studies revealed more recent speciation events in the genus, within the vermilion rockfish and blue rockfish (*S. miniatus* & *S. mystinus*, respectively^17,16^). The two species of blue rockfish were initially named Type 1 and Type 2 and were subsequently described as two distinct species: *S. diaconus (*(Type 1, northern distribution) and *S. mystinus* (Type 2, southern distribution). ^16,18–21^ Burford ^18^ found evidence that the two species diverged approximately 200 kya as a result of geographic isolation. During isolation, both species had large effective population sizes (*N_E_*) and eventually spread into secondary contact in a region covering approximately 420 km coastline of southern Oregon to Northern California, USA.^16^ Across this zone of sympatry, both species co-occur at equal frequencies, yet there was no evidence of hybridization or introgression. However, there was evidence of hybridization in allopatric zones when one species was found in low abundance, suggesting evolutionary reinforcement via prezygotic barriers between these two species in secondary contact.^19^ We hypothesize that this pattern is driven by either ecological segregation or character displacement in the sympatric zone (Fig. 1). A previous study suggested that differentiation in habitat preference (ecological segregation) within the sympatric region could explain species boundaries.^20^ In this region, they suggested that *Sebastes diaconus* inhabits deeper water than in its northern range, whereas *S. mystinus* remains at its typical depth of surface to 25 meters, though ecological segregation as a prezygotic barrier has not been tested. Other abiotic drivers, such as shifts in sea-surface temperature, may also limit species interactions, given evidence that juvenile settlement abundance in both species may be correlated with climatic shifts.^21^

Progress in computational power and molecular methods has increased our ability to measure the effect of natural selection and reinforcement during the speciation process. Reduced representation sequencing such as double digest Restriction-site Associated DNA sequencing (ddRADseq) allows us to produce hundreds to thousands of Single Nucleotide Polymorphisms (SNPs) markers distributed across the genome. These genomic methods enhance our ability to quantify genetic differentiation, detect selective sweeps, and resolve phylogenetic relationships at finer resolutions. In the context of speciation, we can use this approach to conduct an outlier analysis that can reveal genomic regions under directional selection and candidate genes within these regions (e.g. *Aedes*^22^, *Pterois volitans*^23^). However, genome-wide patterns of contemporary differentiation and directional selection have not been evaluated within *Sebastosomus* using next-generation sequencing methods.

To understand the role of secondary contact in reinforcement and investigate evolutionary mechanisms that maintain species boundaries, we analyzed adults and juveniles from allopatric and sympatric regions of the range for both *S. mystinus and S. diaconus* using ddRADseq. Here we address two fundamental questions: (1) is there greater genomic differentiation between the two species in sympatric versus allopatric zones (suggesting character displacement and reproductive reinforcement), and (2) can we identify genomic regions under directional selection within each species when we compare the allopatric with the sympatric parts of their distributions? We also analyzed the degree of geographic genetic differentiation throughout each species’ range to investigate how divergence changes across the geographic gradient shared by both species. Evidence of directional selection in sympatric populations would support reinforcement as a mechanism actively maintaining species boundaries between *S. diaconus* and *S. mystinus*.

## Materials and Methods

### Sampling

We analyzed previously sampled adults and juveniles throughout the distribution of both species (Type 1 *S. diaconus* and Type 2 *S. mystinus*) and focused on regions where the two species are sympatric and allopatric (locations found in Table S1).^16^ The goal was to first confirm differentiation between the two species and then to investigate whether there was evidence of natural selection between allopatric and sympatric portions of each species’ distributions. We first analyzed all locations together to compare genetic patterns between the two species using SNP data, which affords greater power to detect differentiation, to those previously identified using microsatellite and mtDNA control regions analyses.^16,18^ We then tested for greater genomic differentiation between the two species in sympatric versus allopatric populations as a test of genomic character displacement (Question 1). Finally, we addressed questions of directional selection between allopatric and sympatric populations within each species using an outlier analysis (Question 2). We categorized samples in the following ways: *S. diaconus* allopatric, sympatric, or south; *S. mystinus* sympatric or allopatric. The nine individuals of the *S. diaconus* south samples were spread across approximately 310 km at individual locations from Fort Ross to Big Creek (Fig. 1), and represented the only individuals of this species found in these regions. We, therefore, excluded them from the outlier analysis. For the outlier analysis, we compared *S. diaconus* in allopatry versus sympatry and the same for *S. mystinus*.

### DNA extraction, sequencing, and genomic library building

We extracted genomic DNA from fin clips using a Qiagen DNA Extraction Kit (Qiagen Inc., Valencia, California, USA) following the manufacturer protocol. We quantified DNA concentrations using a Qubit fluorometer (Qubit 2.0; Invitrogen, Carlsbad, California, USA). We built ddRADseq libraries using the enzyme pairs *SphI* and *MluCI* and following the protocol outlined in Burford Reiskind et al.^24^ We constructed three libraries for a total of 127 individual rockfish from the two species of rockfish samples (Table S1). We sampled the same individuals from previously published data in Burford & Bernardi ^16^ and Burford,^18^ but we compared allopatric and sympatric populations of each species (Table S1). We conducted single-end sequencing of 100 bp fragments of the libraries on the Illumina HiSeq 2500 following specifications at North Carolina State University Genomic Sequencing Laboratory (Raleigh, North Carolina, USA).

### Bioinformatic pipeline

We used FASTQC (Babraham Bioinformatics; http://www.bioinformatics.babraham.ac.uk/projects/fastqc/) to check the quality of the reads, using a high phred score criterion (phred > 33), prior to processing the barcodes as outlined in Burford Reiskind et al.^24^ We then ran the *process_radtags* script to filter and de-multiplex our variable length barcodes in STACKS v.1.24.^25^ We trimmed the reads to 90 base pairs to make all read lengths identical as required by the STACKS platform. For SNP detection, we ran the *denovo* pipeline (*denovo.pl*) available in STACKS with the following parameters: *m* = 3 (minimum stack depth), *M = 2* (mismatches allowed between loci within an individual), and *n = 2* (mismatches allowed between loci when combining them in a catalog; ^25^). We used the *de novo* pipeline to generate SNP data because there is not a reference genome for either *S. diaconus* or *S. mystinus.* We then used the population pipeline (*populations*) in STACKS with the following parameters: number of populations in which a locus is present (*p*) = 2, proportion of individuals within a population that have these loci (*r*) = 0.75. We chose these parameters to retain loci broadly represented across individuals and populations, following recommendations for *de novo* RADseq analyses in non-model organisms.^26^ We then used PLINK to filter the output from the *populations* pipeline (PLINK v.1.19 http://pngu.mgh.harvard.edu/purcell/plink/) for minimum allele frequency (*maf* = 0.01) and loci with 25% missing data (*geno* = 0.25). While the STACKS pipeline provides the possibility to create datasets in various formats, we used the PLINK format as it is considered more versatile for large NextGen sequenced data. We used the program PGDSpider v.2.1.1.0 (http://www.cmpg.unibe.ch/software/PGDSpider/)^27^ to transform the PLINK dataset to the input file formats required by the software: GENEPOP v.4.2,^28^ LOSITAN,^29^ STRUCTURE v.2.3.4.^30,31^ We recoded our plink files for ADEGENET v 2.1.10 (-- recodeA) to generate input files for ADEGENET in R.^32^

We measured genetic diversity (*H_E_*), inbreeding coefficient (*F_IS_*), and genetic differentiation using pairwise *F_ST_* and Exact test (MCMC parameters: 20,000 dememorization, 500 batches, 10,000 iterations per batch) in GENEPOP. We ran STRUCTURE with 10,000 burnins, 10,000 MCMC replicates, and a *K* ranging from 1 to 6 with 10 iterations per *K* for all species seeded with a user-entered random number. We conducted a second STRUCTURE analysis on the *S. diaconus* population sampled over an expansive geographic area. We used the Evanno Method implemented in StructureHarvester ^33,34^ to determine the likelihood for each value of *K*, assignment of individuals to species, and mixed ancestry of individuals (introgression). The analysis of STRUCTURE on the entire data set confirmed assignment of individuals to species for later outlier analyses. In addition to STRUCTURE, we conducted a Discriminant Analysis of Principal Components (DAPC) in R using the package ADEGENET comparing allopatric and sympatric populations within each species, and also comparing the three *S. diaconus* populations (allopatric, sympatric, and south). We first cross-validated the data sets using 95 replicates to determine the optimal number of principal components (PCs) to evaluate genetic structure. We generated all DAPC and PC figures using the R package *ggplot2*.^35^

To understand the role of selection and local adaptation within each species, we conducted an outlier analysis comparing allopatric and sympatric populations for both species. To identify outlier loci, we conducted an analysis in LOSITAN. We generated outlier loci by first calculating the “neutral” *F_ST_* excluding outlier loci and then ran 10 reps of 1,000,000 simulations in LOSITAN. We applied a false discovery rate (FDR) correction factor of 0.05 (more conservative than the default FDR of 0.10) based on reported high false positive loci rates in LOSITAN’s main algorithm, FDIST2,^36^ and afterwards only accepted outlier loci with *P* – values of 0.001 to reduce false positives.

We compared outlier loci between LOSITAN runs of *S. diaconus* and *S. mystinus* and restricted further analysis to those outlier loci that were shared by both species, to focus on loci that were responding to similar selective force in the sympatric region (see ^37^). We then aligned the sequences containing the outlier loci to the reference genome of *Sebastes entomelas* (GCA_045837885.1; fSebEnt1.0). We used *S. entomelas* reference genome to annotate outlier loci identified through selection analyses, providing a context for regions putatively under directional selection. This two-step approach – *de novo* SNP genotyping followed by *post-hoc* genomic annotation against a closely related reference – is consistent with best practices for non-model organisms where a conspecific reference genome is unavailable.^26^ We aligned outlier sequences to the *S. entomolas* reference using Bowtie2 ^38^ and extracted scaffold coordinates for each alignment using SAMtools.^39^ As the *S. entomelas* genome assembly lacks gene annotation, we extracted the genomic sequence within 15 Kbp of each outlier alignment position using SAMtools faidx ^39^ and submitted these window sequences to the NCBI BLAST nucleotide collection databaseusing megablast.^40^ We considered matches with at least 85% identity and organized by E-value, to identify genes in close physical proximity to outlier loci and thus putatively under directional selection. We further explored these genes using the Zebrafish Information Network ^41^ and NCBI’s Gene database.^42^ We also searched for associated gene ontology (GO) terms including biological processes, molecular functions, or cellular components for each gene symbol through the MGI Batch Query for GO information (http://www.informatics.jax.org/batch) ^43^.

## Results

After filtering first with the *de novo* pipeline then the population pipeline parameters in STACKS, we retained 57,077 polymorphic loci. For analyses within each species, we retained 86,068 polymorphic loci for *S. diaconus* (allopatry, sympatry), 67,542 polymorphic loci for *S. diaconus* (allopatry, sympatry, and south), and 53,787 polymorphic loci for *S. mystinus* (allopatry, sympatry) after filtering. We removed three *S. mystinus* individuals that were missing more than 50% of their data; two individuals in sympatry, and one in allopatry. We did not remove any *S. diaconus* individuals.

### Species classification among marker types

STRUCTURE analysis confirmed the assignment of most of the individuals to the correct species (Fig. 2). Of the 121 individuals also identified using microsatellite and 61 individuals identified using mtDNA sequences in Burford & Bernardi,^16^ we found general agreement with a few exceptions (Table S1). First, when comparing the two marker types for 61 individuals, mtDNA versus nucDNA (microsatellite and SNPs), we found discrepancies in nine individuals in their assignment to species. Four individuals assigned to *S. diaconus* using mtDNA were assigned to *S. mystinus* with the nucDNA markers, all of which were sampled from the sympatric range. Similarly, four individuals assigned to *S. mystinus* in the mtDNA analysis were assigned to *S. diaconus* with the nucDNA markers, including one individual from the sympatric region and three from the northern allopatric zone. In the final comparison between marker types, we found one individual identified as an F_1_ hybrid from mtDNA that was assigned to *S. diaconus* using nucDNA markers (sympatric zone). Comparing the two types of nuclear markers, SNPs versus microsatellites, four individuals out of the 161 did not match. We identified three individuals assigned to *S. diaconus* with the microsatellite data that were assigned to *S. mystinus* with the SNP analysis in this study, one in the sympatric zone and two in the southern allopatric zone. We only found one individual located in the southern allopatric zone assigned to *S. mystinus* with the microsatellite data but was reported as an F_1_ hybrid with the SNP data. This location was outside of the zone where both species are sympatric. For all other individuals, there was agreement among the three marker types, and we used these data to correctly assign and confirm the data set for the outlier analysis (Table S1).

**Fig 2.**
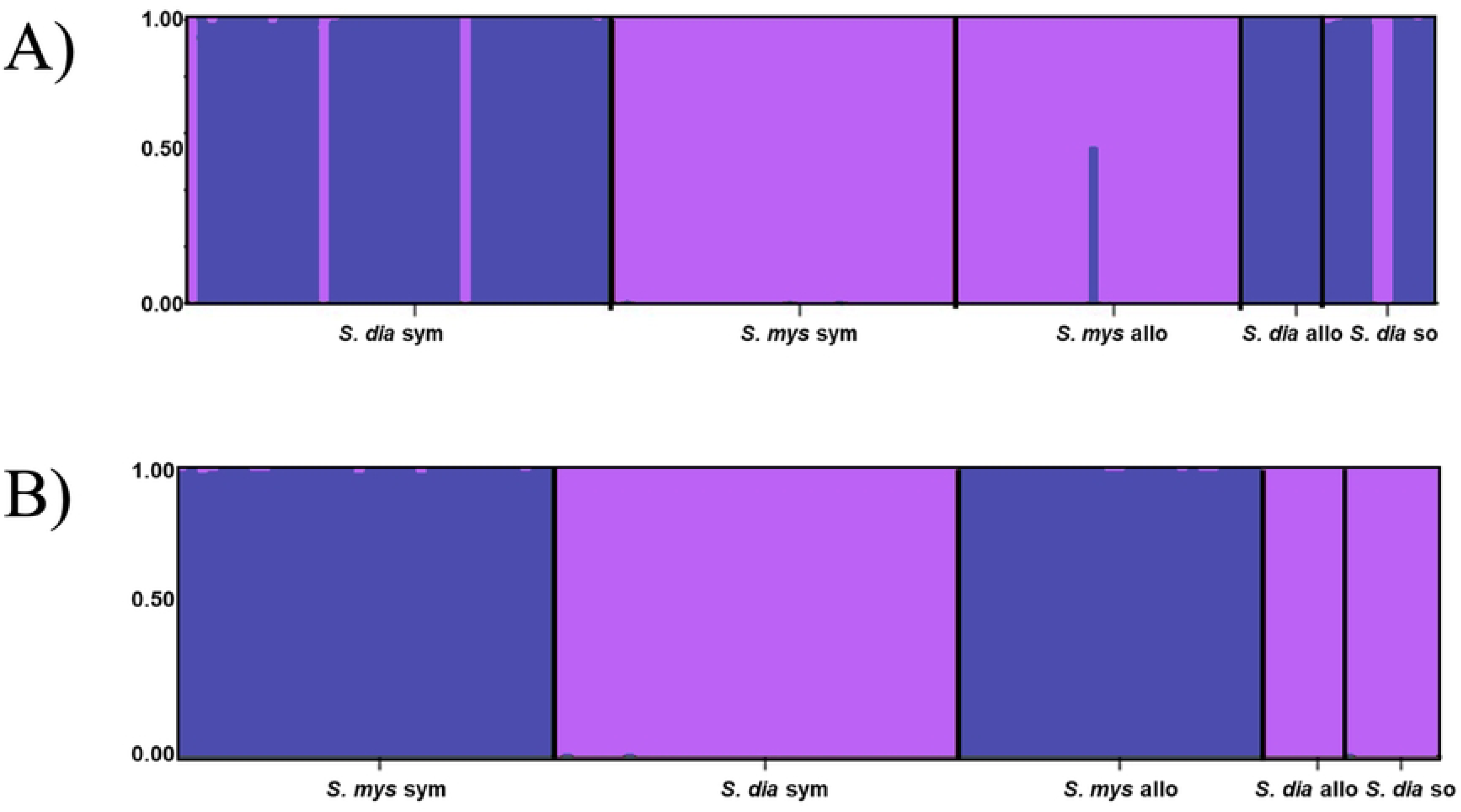
STRUCTURE analysis for both *Sebastes diaconus* and *Sebastes mystinus* throughout both species ranges using 10,000 burnins and MCMC replicates and a *K* (clustering parameter) ranging from 1 to 6 with 10 iterations. We use the Evanno method ^33^ to select the correct cluster of *K* = 2. Here we show the diagnostic analysis (A) and the second analysis with correct species assignment (B) that we use to determine populations in the outlier analysis. Populations include *S. diaconus* allopatric (*S. dia* allo), *S. diaconus* sympatric (*S. dia* sym), *S. diaconus* south (*S. dia* so), *S. mystinus* allopatric (*S. mys* allo), and *S. mystinus* (*S. mys* sym).

### Genetic divergence & genetic relationships between species in sympatry and allopatry

The results of pairwise *F_ST_* values and the Exact test showed significant differentiation when we compared all groups, with the expected significant differentiation between any comparison between the two species (*F_ST_* ranged from 0.331 to 0.338; *P < 0.01*; Table 1). Comparing the two species first in sympatry then in allopatry, we found slightly greater genetic divergence between the species pairs in sympatry (sympatry *F_ST_* = 0.338 versus allopatry *F_ST_*= 0.331; Table 1). We confirmed these patterns in the DAPC analysis comparing sympatry and allopatry between the two species (Fig. 3). We found significant genetic differentiation within the *S. diaconus* species when we compare the northernmost population in allopatry with the samples further south in the *S. diaconus* range (*S. diaconus* allopatry versus *S. diaconus* south; *F_ST_* = 0.007; *P* < 0.01; Table 1). In contrast, we did not detect significant genetic differentiation between *S. mystinus* in sympatry and allopatry.

**Table 1.** Pairwise *F_ST_* values (measure of genetic divergence) among samples of the two blue rockfish species (*S. diaconus* and *S. mystinus*) in sympatry and allopatry across their respective ranges. The comparison is from 57,077 polymorphic SNP loci. The categories are arranged in geographic order from north to south. Statistically significant pairwise comparisons from the Exact test and are bolded (*P* < 0.01 after strict Bonferroni corrections).

|  | *S. diaconus*<br>Allopatry | *S. diaconus*<br>Sympatry | *S. mystinus*<br>Sympatry | *S. mystinus*<br>Allopatry |
| --- | --- | --- | --- | --- |
| *S. diaconus*<br>Sympatry | 0.004 | - |  |  |
| *S. mystinus*<br>Sympatry | **0.332** | **0.338** | - |  |
| *S. mystinus*<br>Allopatry | **0.331** | **0.338** | 0.002 | - |
| *S. diaconus*<br>south | **0.007** | 0.002 | **0.333** | **0.333** |

**Fig 3.**
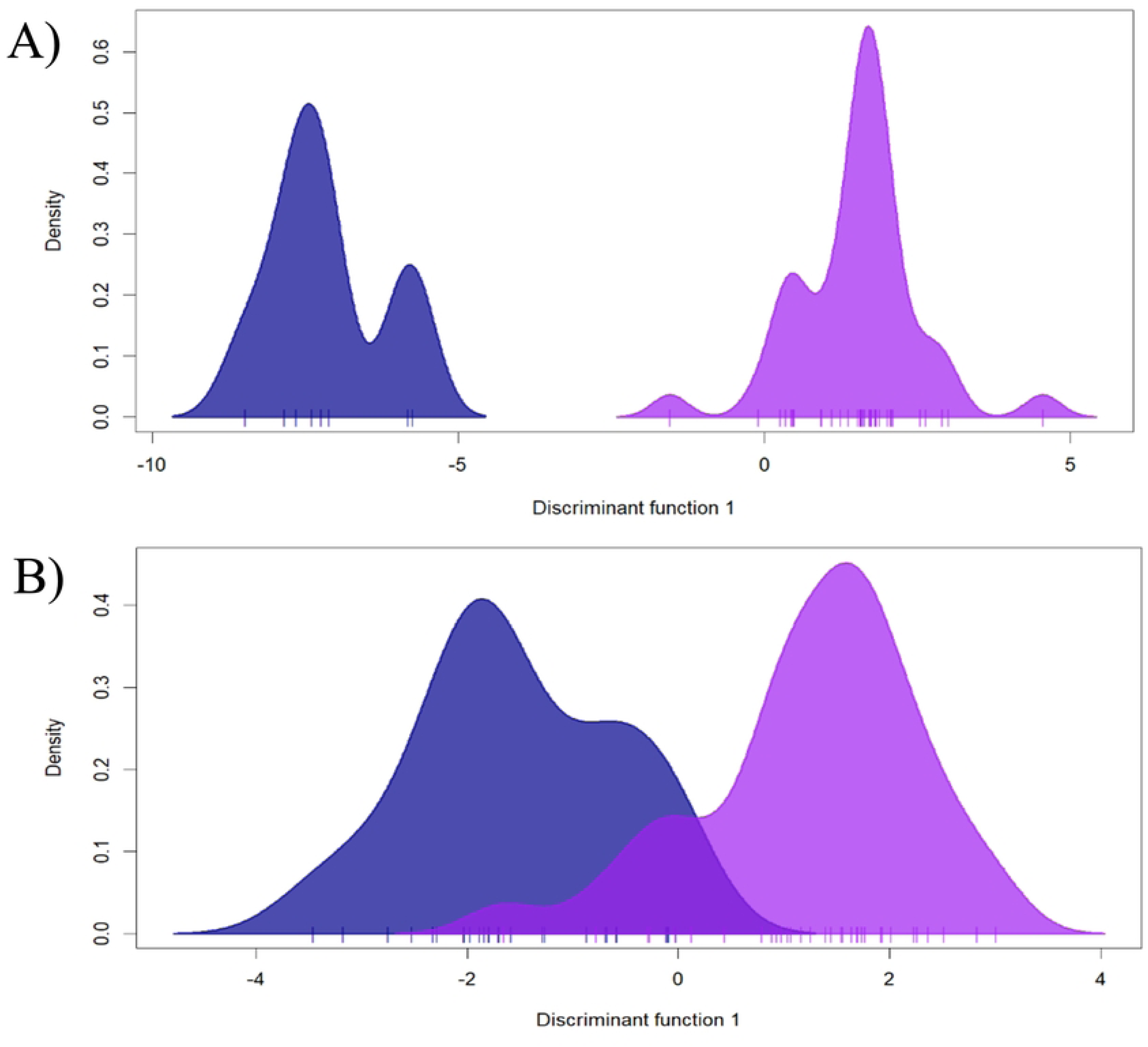
DAPC analysis with 30 principal components comparing allopatric versus sympatric populations for *Sebastes diaconus* (A) and *Sebastes mystinus* (B). We determined the optimal number of principal components by cross-validating the datasets with 95 replicates.

### Genetic characteristics of *S. diaconus*

We used 86,068 loci for *S. diaconus* comparisons between allopatric and sympatric populations. We found that inbreeding was low in both ranges of *S. diaconus*, but slightly higher in the allopatric range compared to the sympatric range (Table 2). The expected heterozygosity was higher than the observed heterozygosity in both the sympatric and allopatric populations (Table 2). We did not find significant pairwise differences between allopatric versus sympatric populations (*S. diaconus* allopatry versus *S. diaconus* sympatry: *F_ST_* = 0.004; *P* > 0.05; Table 1), but found clear separation between allopatric and sympatric populations in the DAPC analysis (Fig. 3A).

**Table 2.** Population genetic summary of *Sebastes diaconus* and *Sebastes mystinus*. This includes observed (*H_O_*) and expected heterozygosity (*H_E_*), and Inbreeding coefficient (*F_IS_*) of both species in allopatry, sympatry and south using 57,077 polymorphic SNPS generated by the ddRADseq analysis.

| Species population | $N$ | $H_O$ | $H_E$ | $F_{IS}$ |
| --- | --- | --- | --- | --- |
| *S. diaconus* Allopatry | 8 | 0.117 | 0.130 | 0.099 |
| *S. diaconus* Sympatry | 42 | 0.120 | 0.129 | 0.067 |
| *S. mystinus* Sympatry | 35 | 0.123 | 0.132 | 0.067 |
| *S. mystinus* Allopatry | 28 | 0.118 | 0.131 | 0.101 |
| *S. diaconus* south | 11 | 0.119 | 0.124 | 0.040 |

The analysis of geographic structure in *S. diaconus* revealed significant genetic differentiation between the northern, allopatric population and the southern population found deep in the *S. mystinus* range. In contrast, the results of the STRUCTURE and DAPC analysis showed that individuals in the sympatric population of *S. diaconus* were genetically similar to both the allopatric and southern populations (Fig. 4). In STRUCTURE, we found support for either a *K = 2* or *K = 3*, depending on the number of replicate runs in the Structure Harvester analysis. Further evaluation of individual assignments supported a *K = 3*, with the sympatric population as intermediate between the two geographically isolated populations in the north and south of this species range (Fig. 4). The sample size of nine individuals in the southern group were excluded from outlier analysis because of their geographic genetic differences and low sample size (Table S1).

**Fig 4.**
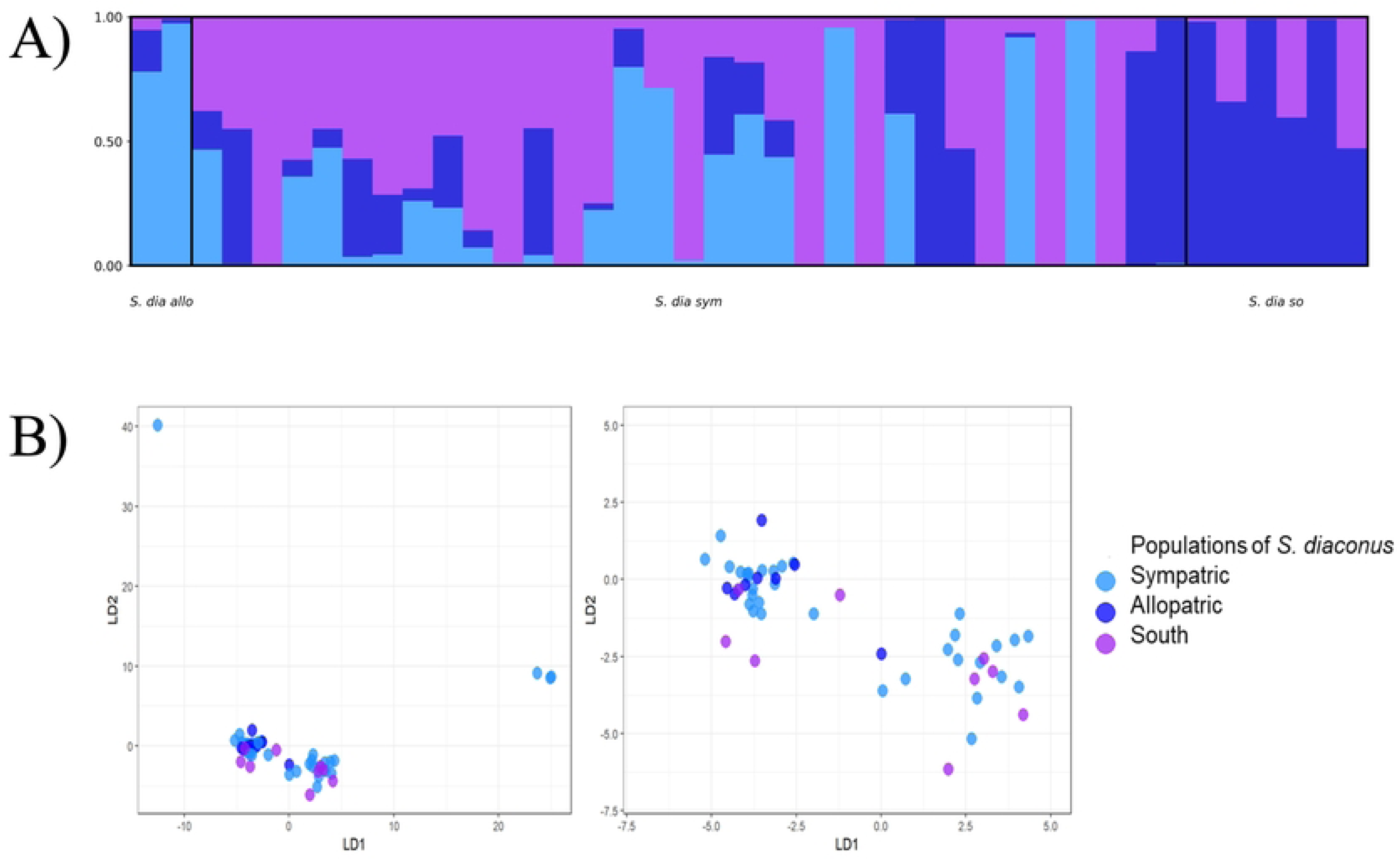
STRUCTURE (A) and DAPC (B) results of the analysis of *Sebastes diaconus* throughout its geographic range. The STRUCTURE results are generated using 10,000 burnins and MCMC replicates and a *K* (clustering parameter) ranging from 1 to 4 with 10 iterations. We use the Evanno method ^33^ to select the correct cluster of *K* = 2 or 3, and we depict the *K* = 3. The DAPC results are generated cross-validated the data sets using 95 replicates to determine the optimal number of principal components (PCs) to evaluate genetic structure. Here we show 30 PCAs for LD1 and LD2 and a zoom in on the most populated area (circle an arrow indicate the region that was zoomed out). Populations include *S. diaconus* allopatric (*S. dia* allo), *S. diaconus* sympatric (*S. dia* sym), and *S. diaconus* south (*S. dia* so).

### Genetic characteristics of *S. mystinus*

To compare the *S. mystinus* allopatric and sympatric populations, we retained a total of 53,787 SNPS. Inbreeding remained low for both populations of *S. mystinus*, with a higher value found in the southern allopatric population than those living in sympatry with *S. diaconus* individuals (Table 2). Measures of expected heterozygosity varied little between the two ranges, but both allopatric and sympatric ranges had observed heterozygosity values that were slightly lower than expected values (Table 2). We did not find significant pairwise difference between *S. mystinus* in allopatric versus sympatric populations (*S. mystinus* allopatric vs *S. mystinus* sympatric *F_ST_*= 0.002; *P* > 0.05; Table 1), and found overlap between the two species in the DAPC analysis (Fig. 3B).

### Outlier loci analysis

We identified 3,348 outlier loci within *S. diaconus* and 840 outlier loci within *S. mystinus* when we compared sympatric and allopatric populations, 61 of which were shared between the two species (Fig. 5). We extracted the sequences of these 61 loci and aligned them to the *S. entomelas* genome (GCA_045837885.1), all of which produced successful alignments. Of 61 outlier loci, 28 (46%) were within 15 Kb of a named candidate gene, six (10%) aligned to regions containing uncharacterized loci with no functional annotation, and 27 (44%) aligned to genomic regions with no gene annotation available in the NCBI nucleotide database.

**Fig 5.**
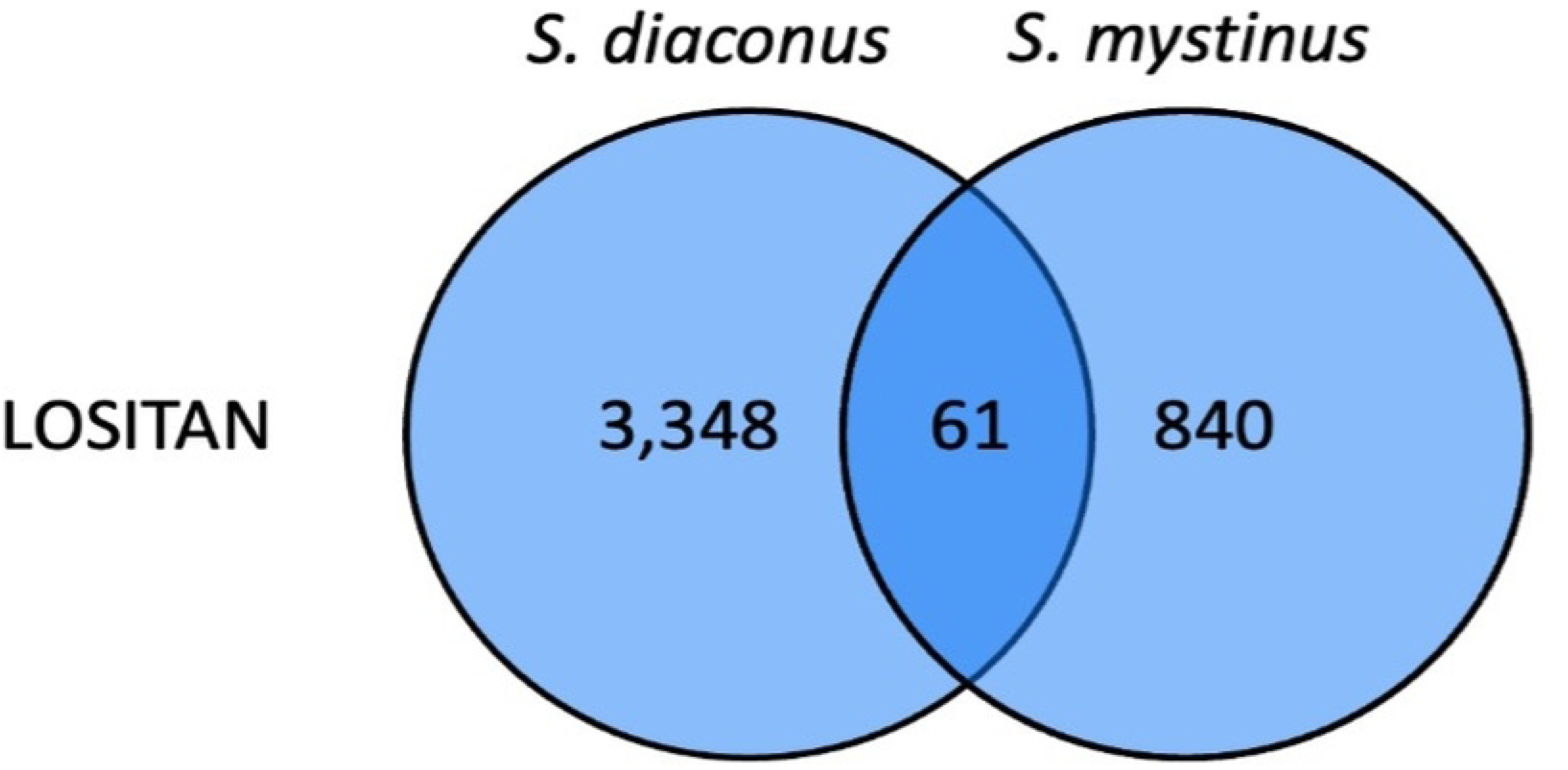
Results of the LOSITAN outlier analysis of allopatric and sympatric populations of *Sebastes diaconus* and *S. mystinus*. Overlapping circles indicated those outlier loci that were shared between the two analyses for each approach. Outlier loci were identified using LOSITAN, first by calculating neutral FST excluding outlier loci, then by running 10 reps of 1,000,000 simulations. We applied a false discovery rate (FDR) correction factor of 0.05 and considered outlier loci with a p-value of <0.001.

The 28 candidate genes fell into several broad functional categories: neurological development and synaptic function (*gad2*, *kcnd1*, *kcnip2*, *srgap3*, *lrrc7*, *cacna1ia*, *mpz*, *pou3f3b*), calcium signaling (*cacna1ia*, *cracr2ab*, *slc24a4b*), sensory perception (*slc24a4b*), transcriptional and epigenetic regulation (*kat6a*, *paip2b*, *wwc1*), cellular stress response and signaling (*mapk11*, *dysf*), and intracellular transport and protein processing (*sar1b*, *qsox1*, *ubl4a*, *lix1l*). GO terms associated with these candidate genes included neurological processes such as GABAergic synapse (GO:0098982), glutamatergic synapse (GO:0098978), postsynaptic density (GO:0014069), voltage-gated calcium channel activity (GO:0005245), and neuronal action potential (GO:0019228), as well as sensory perception terms including phototransduction (GO:0007602), cone photoresponse recovery (GO:0036368), and cone photoreceptor outer segment (GO:0120199; Table S2, S3). The high proportion of unannotated loci (44%) reflects the absence of gene annotation in the *S. entomelas* genome assembly and the phylogenetic distance between *Sebastes* and the most well-annotated fish species in the NCBI nucleotide database, rather than an absence of functional genomic content in these regions.

## Discussion

In this study we confirmed previous findings of genetic divergence between the two recently described species of rockfish, *S. mystinus* and *S. diaconus*, and found evidence that these two species are more genetically diverged in sympatric regions of their distributions compared to allopatric regions. However, the degree of divergence in sympatry was not significantly different in the pairwise *F_ST_* comparison. The lack of significant pairwise differences could be due to biotic interactions with other species in the region diluting the signal between these two species (e.g. ^44^). With greater statistical power of many SNP markers, we were able to better assign individuals to one of the two species throughout their ranges. We also found evidence of greater geographic genetic divergence in the northern species, *S. diaconus*, between the extremes of their range. In contrast, we found no geographic genetic differentiation within *S. mystinus*. While the geographic genetic structure found here differs from previous findings,^21^ the large number of markers identified using ddRADseq provided greater resolution of the geographic genetic structure and species assignment. Finally, we found evidence of directional selection in shared genomic regions of both species, suggesting that selection may keep the two species genetically distinct in regions where both species overlap at higher densities.

Our results confirmed species assignment consistent with previous studies,^16^ though we found greater discordance between nuclear and mitochondrial markers in the sympatric region, suggestive of incomplete lineage sorting. We found further support for lineage sorting by the lack of evidence of hybridization or introgression in any one marker type (Table S1). We did find evidence of at least one F_1_ hybrid juvenile in the allopatric range of *S. mystinus*, which was not found in the previous analyses. Overall, we confirmed the previously identified pattern of introgression between the two species occurred when *S. diaconus* was at lower abundances in the southern part of its distribution.^19^ We found no population genetic structure within the southern distributed *S. mystinus* species, but greater population genetic structure within the northern *S. diaconus* range. This finding was not surprising given the large geographic range covered by *S. diaconus* (greater than 1,600 Km of coastline) and the smaller population size in the southern part of its range. Limited gene flow is expected when the geographic distribution is large, and with smaller population sizes in the south, these populations likely diverged due to genetic drift rather than natural selection.

We did not find the same spatial genetic structure in *S. mystinus* that we observed in *S. diaconus*. The lack of structure within the *S. mystinus* sampling range (greater than 1,200 Km of coastline) suggests larger populations with adequate gene flow along the coastline. Previously, Burford et al.^21^ found evidence of greater geographic genetic structure within *S. mystinus,* possibly the result of species misidentification in the southern part of the range in the previous study.^21^ This suggests that our current analysis was better able to characterize and assign individuals as *S. diaconus* or *S. mystinus* within the southern range of both species, and also better able to diagnose geographic genetic structure. The results from our study provide a clearer picture of range patterns and could suggest that *S. diaconus* is expanding southward, while a similar pattern is not present in *S. mystinus* to the north.

We found almost twice as many outlier loci within *S. diaconus* than we did in *S. mystinus*, suggesting stronger signals of selection within *S. diaconus*. The greater number of outlier loci could be explained by an interaction between gene flow and selection (see ^45^). However, given the geographic genetic structure described above, some of these outlier loci are likely due to genetic drift or limited gene flow rather than selection among the populations of *S. diaconus*. Given that genetic drift or limited gene flow was an important contributor to genetic divergence in *S. diaconus*, we restricted the outlier analysis to loci shared by both species. We recognize that restricting to shared outlier loci between the two species is conservative, because it could eliminate species-specific genomic regions where selective forces, either similar or different, influence different genes within each species. However, given our goal of finding outlier loci within or near genomic regions under selection, a more conservative approach was appropriate. The outlier analysis showed evidence of shared genomic regions of directional selection in *S. mystinus* and *S. diaconus*. The fact that these regions are shared between species suggests they may be responding to similar selective pressures, whether driven by abiotic environment or biotic species interactions. Several traits could allow these two species to coexist in sympatry with little hybridization, including temporal or spatial separation or divergence in reproductive timing. For example, there is evidence that depth partitioning maintains species boundaries of the vermilion rockfish (*S. miniatus*)^17^ and this was also suggested as a mechanism for physical separation of *S. diaconu*s and *S. mystinus* in the overlapping range.^20^ To a lesser extent, microgeographic partitioning within the reef could maintain the barriers between *S. mystinus* and *S. diaconus*.^19^ A third possibility is temporal separation due to differences in reproductive timing between these two species, a common characteristic found between many species in the *Sebastes* genus (e.g. ^46,19^). Alternatively, there could be changes in mating displays or reproductive morphology that could be in place prior to secondary contact, but weaken when one species is at lower densities,^18^ potentially explaining the observed introgression in the southern region. Finally, both species may be adapting to similar environmental pressures in their shared habitat, which could explain the overlap in genomic regions under directional selection. This means that both species are influenced by interactions with their abiotic environment in sympatry rather than interspecific interactions. To better discern the driver of reinforcement against introgression in sympatry requires further understanding of these shared genomic regions under directional selection.

Our BLAST results identified putative genes with functions supporting phenotypes that might maintain species boundaries in regions of overlap. We found the highest number of genes under selection related to neurological function, specifically genes involved in synaptic transmission and postsynaptic organization (*gad2*, *srgap3*, *lrrc7*), ion channel activity (*kcnd1*, *kcnip2*, *cacna1ia*), neural development and myelination (*pou3f3b*, *mpz*, *ndrg1*). The enrichment of outlier loci near genes involved in synaptic transmission and neural development is consistent with findings in other fish undergoing speciation, including within the genus *Sebastes*. In depth-segregated *Sebastes* species pairs (*S. chlorostictus* – *S. rosenblatti* and *S. crocotulus* – *S. miniatus*), genomic islands of divergence are enriched for neurosensory genes, suggesting a role for neural gene divergence in early *Sebastes* speciation.^47,48^ Similar patterns have been observed in other teleosts such as lake whitefish, where genomic regions under divergent selection are enriched for neurological and behavioral functions.^49^ More broadly, neurogenomic divergence at synaptic transmission genes has been linked to speciation in other vertebrate systems. For example, in chorus frogs, seven synaptic transmission genes diverged between sympatric and allopatric populations, with gene networks enriched for neurotransmission showing the strongest divergence.^50^ These findings, coupled with the neurological and myelination genes identified here, suggest that divergence in neural signaling pathways may be a general mechanism by which reinforcement maintains species boundaries in secondary contact zones.

We found one gene, solute carrier family 24 member 4b (*slc24a4b*), which is expressed in the eye.^51^ *Slc24a4b* is associated with cone photoreceptors and assists with adjustment to various levels of lighting in mice and zebrafish.^51^ This could suggest directional selection in eye development related depth differences. Sivasundar and Palumbi ^52^ also found evidence for positive selection in rhodopsin associated genes in rockfish species that occupy different depths in the water column. Their results are similar to the associations found with Ornithine decarboxylase-like (*Odc1*) in rockfish species by Behrens et al. This finding further supports the evidence that depth is a prominent driver of speciation in *Sebastes*.^47,48^

Several outlier loci were also associated with genes involved in transcriptional and epigenetic regulation, including *kat6a*, a histone acetyltransferase involved in chromatin remodeling and developmental regulation, and *pou3f3b*, a transcription factor with roles in forebrain development. Divergence at regulatory loci may suggest differences in gene expression may also contribute to reinforcement of species boundaries in sympatric zones.

The functional diversity of candidate genes identified among outlier loci suggests that reinforcement of species boundaries in *S. diaconus* and *S. mystinus* is likely polygenic and involves divergence across multiple biological pathways. While the functional roles of many outlier loci remain unresolved, the identification of neural and regulatory gene candidates support patterns observed in other fish speciation systems and support a role for neurogenomic divergence in driving speciation within sympatric zones.

The use of a de novo pipeline for SNP genotyping in this study warrants explicit consideration, as reference-guided approaches are increasingly common where suitable genomes exist.^53,54^ No reference genome is available for either *S. diaconus* or *S. mystinus*. The closest relative with a published genome, *S. entomelas*, is sister to the *S. diaconus* and *S. mystinus* clade within the subgenus *Sebastomas*, with divergence estimated to have occurred between 3 and 5 mya. ^55,56^ Aligning short reads to a heterospecific reference at this level of divergence introduces ascertainment bias through non-random locus dropout.^57–59^ This dropout is most severe at regions most likely to harbor signals of selection and reproductive isolation, which are the focus of this study. Furthermore, Paris et al. explicitly demonstrated that building loci *de novo* and subsequently integrating reference alignment positions outperforms aligning raw reads directly to a reference genome, supporting the two-step approach employed here: *de novo* SNP genotyping followed by post-hoc annotation of outlier loci against the *S. entomelas* genome. ^26^

## Conclusion

In this study we have further categorized the population dynamics of these two species and highlighted genomic regions and putative genes of interest for future studies. Future research should focus on the regions of secondary contact, characterizing variation in these candidate genomic regions to validate signatures of selection. Finding and validating evidence of selection will allow us to begin to understand how these two sister species avoid hybridization when occupying the same geographic areas at similar densities. Combined with observational and descriptive data on the ecological and demographic differences between the two species (e.g., reproductive timing, juvenile recruitment timing and location, adult depth and feeding differences) will advance our understanding of both the speciation process and reinforcement that has maintained species boundaries. Elucidating the reproductive barriers for these recently described rockfish species will contribute to conservation management strategies for both of these species that are part of active fisheries in the eastern Pacific.

## Acknowledgements

We would like Kara Carlson and Megan Dillon for their input on the original study and manuscript and Michael Hay Reiskind for logistical support for the genomic methods. We thank Emily M.X. Reed for her assistance in training authors on wet laboratory procedures used in this study. We would like to thank the GG Scholars graduate training program as part of the NC State University Genetics and Genomics Academy.

## Supporting information

**S1 Table. Results of the comparison of individual assignment and finalized comparisons for the outlier analysis of *Sebastes mystinus* and *Sebastes diaconus* in sympatric and allopatric sampling zones.** Individuals are arranged based on preliminary Structure analysis using 57,077 loci generated in the ddRADseq analysis. The finalized assignment is used for the pairwise *F_ST_* and outlier analysis and compared to previously published data from Burford & Bernardi (2008)’s microsatellite data and mtDNA Control Region sequence data.

**S2 Table. Results from NCBI Nucleotide BLAST search of exons within 15 Kb of outlier loci shared by *Sebastes diaconus* and *Sebastes mystinus*.** We report results from the BLAST for sequences that had a percent identity of at least 85% and with the lowest E-value. No Matches indicates no gene hits within the search results, and No Significant Matches indicates sequences that generated numerous gene results with low query coverage as we could not differentiate the matching gene.

**S3 Table. GO term results from the MGI database for candidate genes shared by *Sebastes diaconus* and *Sebastes mystinus* found in NCBI BLAST search.** We include candidate genes found via BLAST (Table S2) into a batch query to gather GO terms.

## Notes

### Competing Interest Statement

The authors have declared no competing interest.

